# Building predictive Markov State Models of ion channel permeation from Molecular Dynamics

**DOI:** 10.1101/2024.02.22.581422

**Authors:** Luigi Catacuzzeno, Maria Vittoria Leonardi, Fabio Franciolini, Carmen Domene, Antonio Michelucci, Simone Furini

## Abstract

Molecular dynamics (MD) simulation of biological processes has always been a very challenging task due to the long timescales of the processes involved and the challenges associated with handling the large amount of output data. Markov State Models (MSMs) have been recently introduced as a powerful tool in this area of research, as they provide a mechanistically comprehensible synthesis of the large amount of MD data and, at the same time, can be used to estimate experimental properties of biological processes. Of the many studies on protein simulation and the MSM-assisted approach, only a few have addressed ion channel permeation and, more importantly, none of these have tried to build a model capable to predict the currents passing through the channels, which are ultimately crucial for comparing simulations with experimental results. Herein, we propose a method for building an MSM of ion channel permeation that correctly evaluates the current flowing through the channel. This was done by including in the model the definition of a flux matrix carrying information on the charge moving across the channel, suitably built to be used in conjunction with the transition matrix to predict the ion current. The proposed method is also able to drastically reduce the number of states so to obtain an MSM simple enough to be easily understood. Finally, we applied the method to the KcsA channel, obtaining a four-state MSM capable of accurately reproducing the single channel ion current from microseconds MD trajectories.

## Introduction

Ion channels are membrane proteins that allow the passage of ions across cell membranes, remarkably combining high throughput rate and high selectivity, and this combination of properties has interrogated scientists for decades. A turning point in the understanding of these extraordinary features has been the possibility to obtain high resolution atomic structures of ion channels, fundamental to relate the permeation and selectivity processes to the chemical structures forming the ion channel pore (1). Among the advantages offered by the knowledge of the ion channel structure, there is the possibility to perform Molecular Dynamics (MD) simulations (2, 3).

All-atom MD simulations, which defines the state of the system by considering the coordinates and velocities of all atoms, consist of a numerical integration of the classical equation of motion, combined with algorithms for sampling configurations in the proper thermodynamic ensemble. Because of this complexity, at present one microsecond MD simulation – a time window that allows to observe only a scanty number of ions crossing a 20 pS-conductance channel at 100 mV – requires weeks of ordinary workstations workout. This limited amount of information is not sufficient to study and understand mechanisms such as permeation and selectivity. A second major MD simulation shortcoming is the overwhelming amount of data to process (i.e. the positions and the velocities of all the atoms over time), most of which are irrelevant for the understanding of the processes of interest. This massive amount of information might hinder the identification of forces and structures relevant for the process under investigation. Given all the above, other strategies should be considered to complement MD simulations to appropriately address permeation in ion channels. One interesting option is to integrate a Markov State Model (MSM) approach to MD simulations.

MSMs are emerging as powerful strategies for extracting relevant information from MD trajectories in a statistically robust way (4). The analysis of MD trajectories using MSMs entails the following steps (5): (i) the definition of a preliminary set of input properties; (ii) the clustering of the trajectories using these input properties; (iii) the calculation of the transition matrix among these clusters; and (iv) the coarse graining of the estimated MSM into a limited number of long-lived metastable states. These metastable states group together microscopic states that rapidly interconvert into each other, while separating microscopic states that are distant in time, offering in this way a simplified description that captures the most important dynamic events in the system. Previous applications of MSMs to the analysis of permeation and selectivity in ion channels have been reported by Harrigan et al., 2017 (6), Domene et al., 2021 (7) and Lam and DeGroot 2023 (8). These studies provide a proof-of-principle of how MSMs can be useful for the analysis of permeation and selectivity, but do not provide a formalism to predict the ion channel current from the MSM, a necessary step to compare the model’s results to experiments.

The purpose of this study is to build a method that correctly recovers from MD data an MSM of ion permeation, defined by a limited number of discrete states and the corresponding transition rates among these finite states. Such an MSM offers an immediate understanding of the system dynamics. We will also test the method by applying it on MD data obtained with the bacterial KcsA potassium channel (9). This channel has similar structural properties to many eukaryotic K channels, and because of this it is widely studied to understand K transport (10, 11). KcsA is formed by the juxtaposition of four identical subunits, each composed of two transmembrane segments, TM1 and TM2, that make up the permeation structure (Figure 1A, left). At the pore extracellular entrance, the four P-loops connecting the four TM1 and TM2 segments of each protein chain make the narrow selectivity filter (SF), about 12 Å long and 3 Å wide, formed by a highly conserved sequence of amino acids (TVGYG in most K channels), with their carbonyl oxygen atoms pointing towards the pore. Together with the hydroxyl oxygens of the threonine residues, these carbonyl oxygens define a series of binding sites for K, named S1 to S4 (Figure 1A, right). Two additional external binding sites, immediately at the intracellular and extracellular entrances of the SF, called S0 and S5, were also identified (11–13). We performed extensive MD simulation by obtaining tens of K ion passages through the SF. We then used these data to build a reduced MSM containing only four relevant states and capable to reproduce an ion current comparable to that obtained by directly counting the ion passage events from MD data.

**Figure 1.**
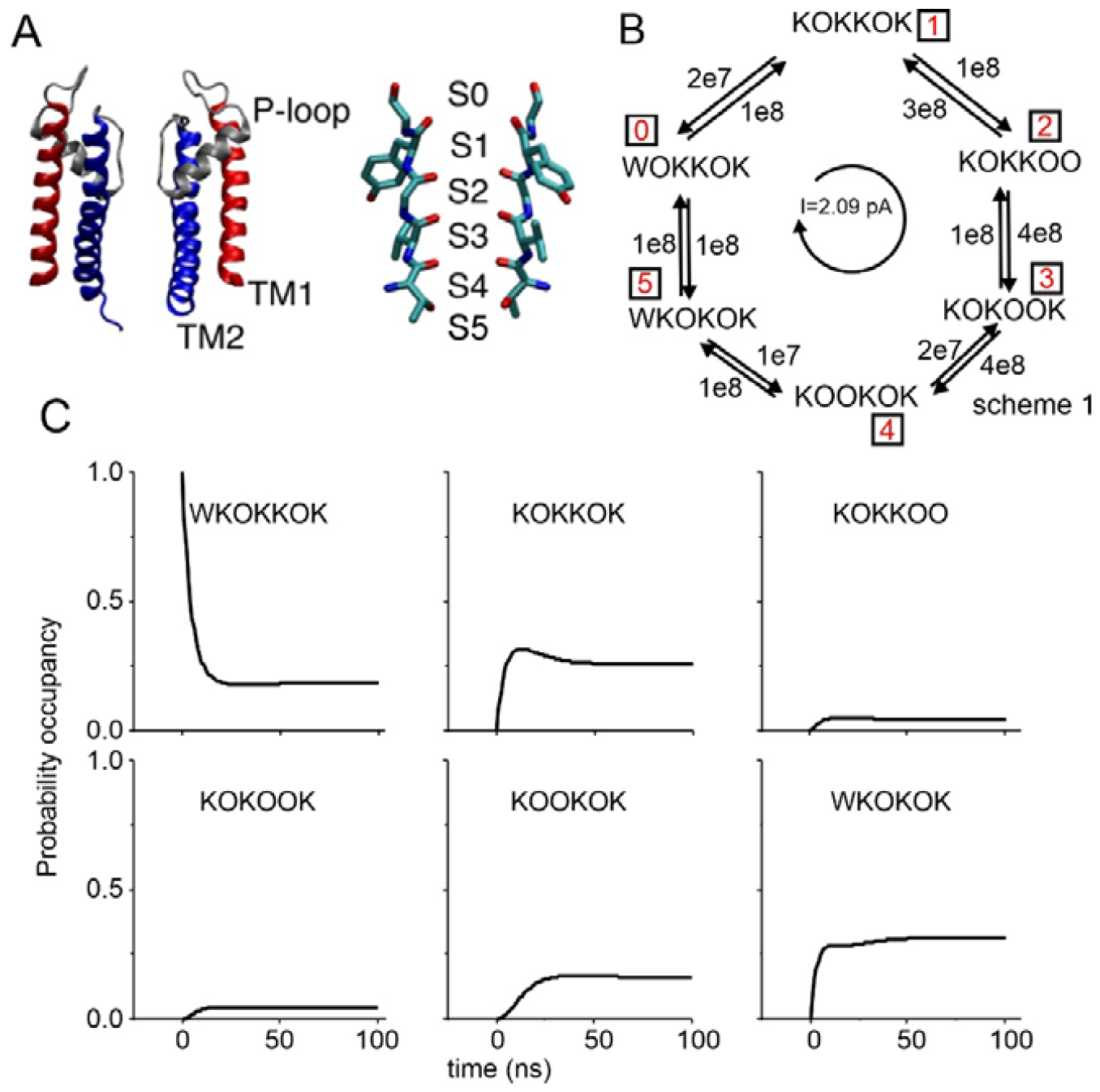
**A)** Atomic structure of the KcsA channel (left, only two of the four subunits are shown) and its SF (right), showing the six binding sites (S0-S5). **B)** Exemplificative 6 states MSM considered in the text. States 0 to 5 correspond to different occupancy states of the SF. **C)** Prediction of the occupancy of the six states of the scheme in B vs time (eqn. 3), starting from an initial condition in which the probability occupancy of state 0 (WOKKOK) is unitary.

## Materials and Methods

### Molecular Dynamics (MD) simulations

The channel model was based on the experimental structure of KcsA-E71A in the open/conductive state, Protein Data Bank entry 5VK6. The entire transmembrane domain of the channel, from residue Trp26 to residue Gln121, was considered. The initial atomic coordinates of the systems were obtained with CHARMM-GUI. The lipid membrane was a mixture of 1-palmitoyl-2-oleoyl-glycero-3-phosphocholine (POPC) and 1-palmitoyl-2-oleoyl-sn-glycero-3-phosphate (POPA), with a ratio of 3-POPC/1-POPA. The channel was inserted in the lipid membrane as defined in the Orientations of Proteins in Membranes (OPM) database. The system was solvated using TIP3P water molecules (∼15,000 molecules) and ions to reach 200 mM KCl. Potassium ions were manually placed at binding sites S0, S2, and S4. The ff14sb version was used, in combination with ion parameters by Joung and Cheatham for the TIP3P water model. VdW interactions were truncated at 9 Å. Standard AMBER scaling of 1−4 interactions was applied. The equilibration protocol consisted of 10,000 energy minimizations, followed by 10 ns in the NPT ensemble with timesteps equal to 1 fs and 60 ns in the NPT ensemble with a timestep equal to 2 fs. During the equilibration protocol, restraints on protein and lipid atoms were gradually reduced to zero. Long-range electrostatic interactions were calculated with the particle mesh Ewald method using a grid spacing of 1.0 Å. The SETTLE algorithm was used to restrain bonds with the hydrogen atom. The temperature was controlled at 310 K by coupling to a Langevin thermostat with a damping coefficient of 1 ps−1. A pressure of 1 atm was maintained by coupling the system to a Nose−Hoover Langevin piston, with a damping constant of 25 ps and a period of 50 ps. The presence of membrane potentials was mimicked by applying a constant electric field acting in the direction perpendicular to the lipid membrane. The simulations with the external electric field were performed in the NVT ensemble. NAMD2.11 was used for all the simulations.

### Transition matrix from MD trajectories

Trajectories were analyzed using the python library MD analysis and the SciPy ecosystem. Visual Molecular Dynamics (VMD) was used to inspect trajectories and to generate images of the systems. The center of mass of the following set of atoms were considered as boundaries for the different channel regions: residues Ala107, hydroxyl oxygen atom of residues Thr75, backbone oxygen atom of residues Thr75, Val76, Gly77, Tyr78 and Gly79. Ions were placed in binding sites S0-S4 or in the cavity when in-between the boundaries of the corresponding binding site, and when within 4 Å from the channel axis (8 Å for ions in S5). Using these binding site definitions, each frame of the simulated trajectories was converted into a string describing the occupancy for a posteriori classification. The occupancy state of the channel in every frame of each trajectory was case coded using the following regular expression: [Kw][Kw-][Kw-][Kw-][Kw-][Kw-]. The first character of the string was K when a potassium ion was in S5 or w if the cavity was only filled with water molecules. Similarly, the next character of the string was K if S4 was occupied by an ion, w if it was occupied by a water molecule, and - if S4 was vacant. To define the characters describing binding sites S3 to S0, the same criteria were imposed. At times, due to the intrinsic structural nature of S0, an ion and a water molecule can coexist at S0, and thus, in this case, the character K was selected. This classification of the occupancy state of the channel was used to convert the simulated trajectories into sequences of discrete states to be used to estimate the transition matrixes count state-to-state transition with the sliding method.

## Theory

### A Markov State Model formalism of ion channel permeation

MSM have been used for more than 50 years to describe the ion permeation through biological ion channels (14), thus long before the knowledge of their atomic structure. In all these models a certain number of configuration states are identified, each one representing a different arrangement of permeating ions bound to the different binding sites along the permeation pathway (i.e. the channel pore). The tendency of the system to transit from a configuration state, *i*, to another one, *j*, is defined either in terms of a transition rate constant, Q_ij_, or in terms of the probability that the *i* → *j* transition occurs in a certain time interval or lagtime Δt, T_ij_(Δt), both constants in time in a homogeneous MSM (an MSM is defined homogeneous when the probability of going from a state *i* to a state *j* only depends on the lagtime, but not on the absolute time). Q_ij_ and T_ij_(Δt) are related to each other by the following relationship:

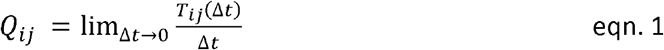

Configuration states and transition rate constants are usually sketched with kinetic schemes (as used in chemical reactions), with arrows between the configuration states indicating possible transitions, and numbers associated to each arrow indicating the numerical value of the transition rate constants. For exemplificative purposes, we will consider a generic MSM of K permeation through a K-channel, characterized by the kinetic scheme in Figure 1B. The scheme is inspired by the prokaryotic K-channel KcsA (Figure 1A). Accordingly, the model considers six adjacent K binding sites along the channel pore (going from the outside to the inside, called S0 to S5). At any given time, each site can be empty (indicated with ‘O’), bound to a K (‘K’), or bound to a water molecule (‘W’). In the simple example, only six states, among the many possible ones, are considered to be possible.

#### Q and T matrices of MSMs

The kinetic rate constants of an MSM can also be grouped in a Q matrix, where the element Q_ij_ represents the kinetic rate constant going from state *i* to state *j*, and the diagonal elements Q_ii_ are defined as 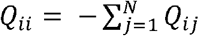, with *i*≠*j*, where N is the total number of states (6 in our example; the reason for the above definition is that it make possible a simple relationship with the transition probabilities, see below). For the Scheme of Figure 1B, the Q matrix is:

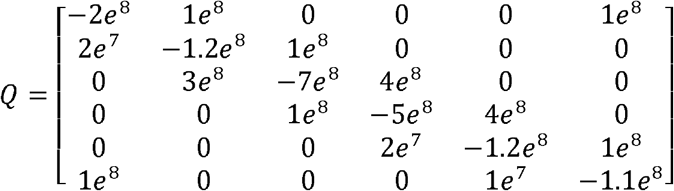

where rows (columns) 0, 1, ….5 correspond to states WOKKOK, KOKKOK, ……WKOKOK.

Similarly, the transition probabilities for a certain lagtime (Δt) can be represented as a transition probability matrix T(Δt), where each element T_ij_(Δt) represents the probability of the system to go from a state *i* to a state *j* in a time Δt, given that the system was in state *i* at time zero. In the T(Δt) matrix, the diagonal elements are the probabilities that no transition occurs in the lagtime Δt, so that 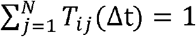 for all possible *i*. It can be shown (see appendix A) that for homogeneous MSMs the T matrix is related to the Q matrix by the following relationship:

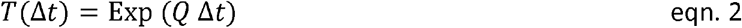

where Exp represents the matrix exponential operator and Δ*t* the lagtime. Notice that, in contrast to eqn. 1 that is valid only for very small lagtimes, eqn. 2 is correct for any chosen lagtime.

Knowing the transition probability matrix at a certain lagtime Δt for an MSM allows to predict the occupancy of the various states of the model at successive times. More specifically, if the fractional occupancy of the system at a certain time t is represented by the row vector P(t)=[p_0_(t) p_1_(t) p_N_(t)], where p_i_(t) represents the probability of finding the system in state *i*, then:

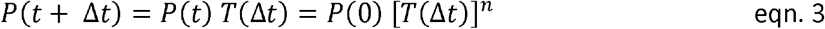

where n is chosen so that t+*t* =n Δ*t*. A similar equation may also be written for the time evolution of the transition probability matrix:

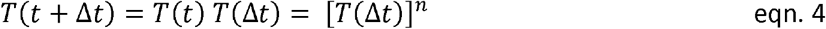

The above equations, known as different forms of the Chapman-Kolmogorov equation, clarify a fundamental property of homogeneous MSMs: the occupancy of the system at a certain time only depends on the status of the system at the previous time step, but not on its older history. Figure 1C shows the evolution of the occupancies of the 6 states of our illustrative MSM, predicted by eqn. 3, taking a lagtime of 1 ns and assuming that the occupancy of state 0 (WOKKOK) is 1 at time zero.

The transition probability matrix of an MSM has the useful property that if an eigen-decomposition of its transpose is performed, the eigenvector corresponding to the unitary eigenvalue (always present) gives the equilibrium fractional occupancy of the various states, expressed as a row vector, *P*_*inf*_. For our illustrative MSM, performing the described operation we obtain:

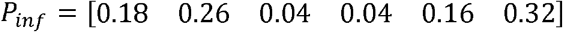

Figure 1C shows that the equilibrium condition is reached after about 50 ns.

#### The flux matrix of MSMs

The kinetic scheme of an ion permeation model does not fully define its behavior. To predict the ion current, that is, the ion channel permeation observable during experimental recordings, it is necessary to also know the amount of charge transferred through the permeation pore during each transition. Assuming that the movement of a K from a site to the next one immediately to the right (thus, towards the outside; Figure 1A) has a +1/7 value (if the complete passage of a K from the internal to the external solution has a value of +1), the flux matrix for our MSM example, that is the matrix reporting the charge flux associated to each possible transition, will be that shown in Figure 2B.

**Figure 2.**
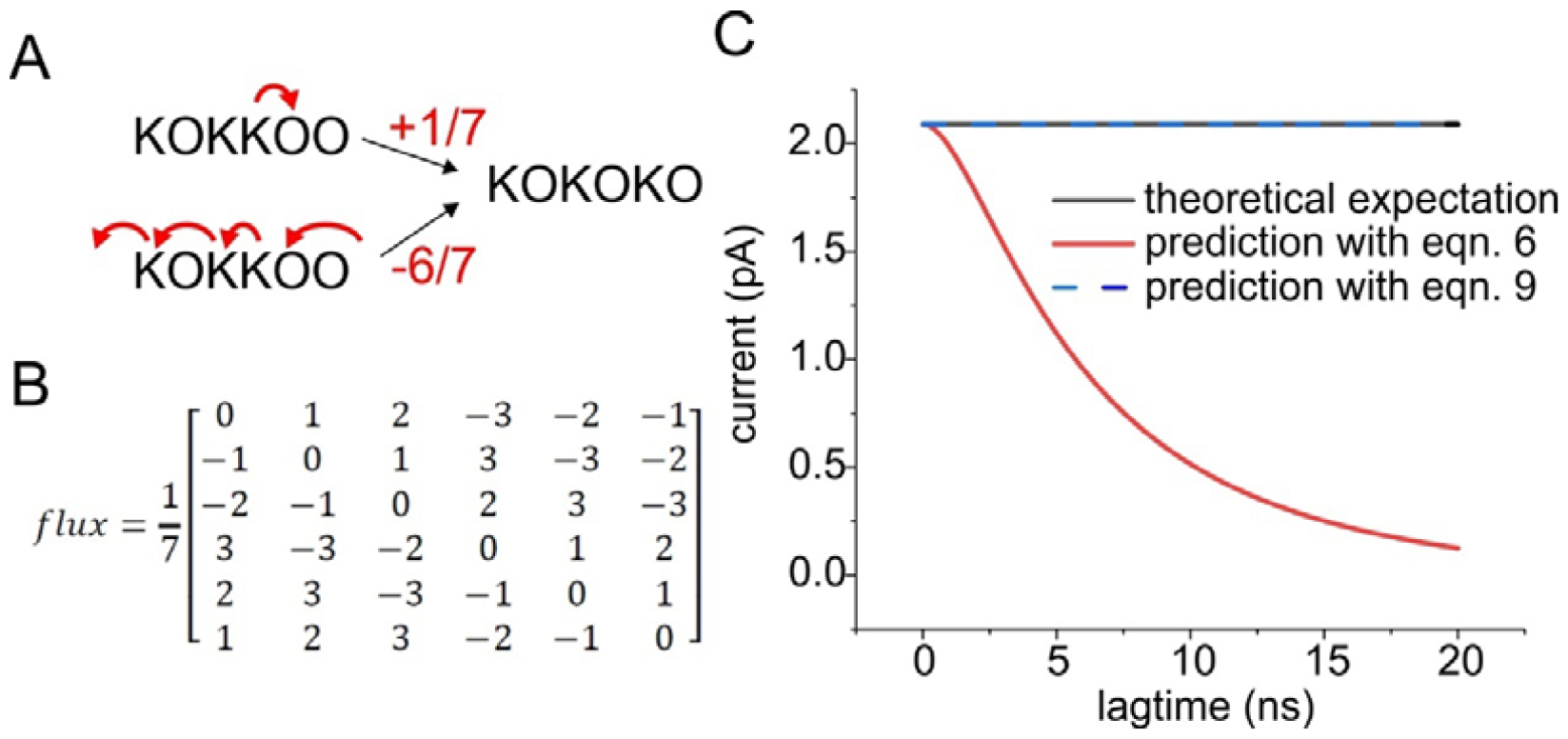
**A)** Amount of charge translocated according to the route taken to make the transition. **B)** Flux matrix for the scheme in Fig. 1B. **C)** Prediction of the ion current using transition matrices at varying lagtimes for the 6-state MSM, using eqn. 5 (black), eqn. 6 (red) and eqn. 9 (blue). Note that the theoretical expectation (black) superimposes exactly with eqn. 9 prediction (blue)

Such a matrix can be computed automatically using the algorithm reported in Appendix B. Notice that there are, in general, multiple possibilities in going from *i* to *j*, and the algorithm of Appendix B selects the pathway reached with the minimum number of transitions. For example, the transition KOKKOO → KOKOKO have an associated flux of +1/7, obtained by moving the K sitting in S2 to S1. It is however possible to reach the same final configuration by moving K ions in the reverse direction, letting the K ion in S5 to exit in the internal solution, moving the K ion from S3 to S5, the K ion from S2 to S3 and an outside K ion to S1, giving an overall flux of -6/7 (Figure 2A). The first transition is that chosen for the flux matrix since it is obtained only in 1 step instead of the 6 steps needed for the second transition.

#### Assessing ion currents from MSMs

There are several ways to assess the ion current (I) produced at steady state by an MSM of ion permeation. The most straightforward is to assess the net rate of entrance (exit) of a K ion from the inside (to the outside). For our exemplificative scheme of Figure 1B this would give:

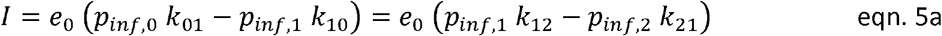

Where *e*_0_ represents the elementary charge, carried by a K ion. A more general expression, working also in more complicated MSM showing multiple points of entrance/exit of ions, and using both the Q matrix and the flux matrix, is the following:

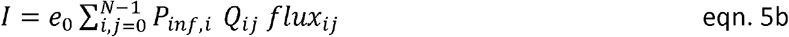

In our example, both the above equations 5 gave a current of 2.09 pA. An alternative to eqn. 5b to assess the ion current associated to an MSM is eqn. 6 that uses information from the transition probability matrix instead of the Q matrix, as suggested by eqn. 1.

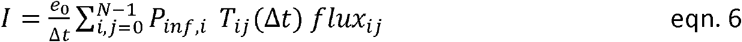

As shown by the red line in Figure 2C, that tracks the output current given by eqn. 6, the estimated current depends greatly on the lagtime used, predicting the correct current at infinitesimal short lagtimes, but departing markedly from the theoretical expectation at larger lagtime values of transition probability matrices. The reason for missing the correct current prediction when using transition probability matrices obtained with large lagtimes originates from the fact that, of the many possible pathways, the flux matrix used considers the charge carried following the pathway with the minimum number of steps. While this approximation may be valid for very short lagtimes, increasing the interval, multistep pathways become more and more likely and, therefore, it is necessary to consider them for a correct prediction. This means that, in general, we need to use a composite flux matrix that considers the charge carried along all possible pathways that might be followed. More specifically, from eqn. 4 the transition probability from state *i* to state *j* may be written as

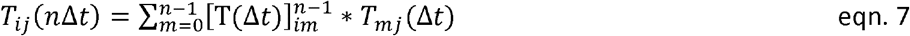

that decomposes the probability of going from state *i* to state *j* into the different pathways that can be followed. Using the above property, we can assess the composite flux matrix F valid for a lagtime of nΔt using the following recursion:

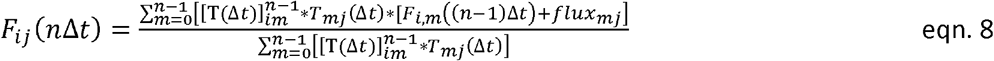

starting from the flux matrix calculated at the shortest lagtime available using the algorithm described in Appendix B. The F matrix obtained in this way will contain the average charge carried along each transition, obtained by a probability weighted average of all possible pathways, and can be used to predict the current in association with the transition probability matrix obtained for the same lagtime using the following relationship:

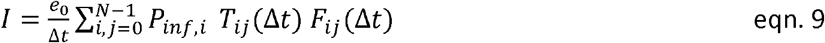

The blue line in Figure 2C shows that eqn. 8 and 9 predicts the correct value of the ion current using transition probability matrices obtained at every lagtime.

In conclusion, a homogeneous MSM of ion permeation is fully defined by its transition probability matrix obtained at a relatively short lagtime T(Δt), and the corresponding flux matrix ‘*flux*’ that can be assessed computationally. With these two pieces of information, we can predict the ion current. A correct prediction of the ion current can also be recovered using transition matrices obtained at larger lagtimes nΔt by finding a corresponding composite flux matrix F(nΔt) that considers the average charge flux carried during nΔt.

### Markov state model reduction and ion current assessment for the resulting quasi-Markov state model

A distinctive feature of MSM built from MD data is the large number of metastable states that needs to be considered. For example, in the case of KcsA channel that has 6 binding sites where K, water, both, or none of them can be present, a total of 4^6^=4096 possible states should be considered (7). Obviously, to have an MSM that could help to understand the mechanism of ion permeation, one may need to greatly reduce the number of possible states, using some reasonable reduction strategy that lump together states.

#### MSM reduction strategy

We next illustrate a possible strategy for MSM reduction that preserves steady state occupancy and flux rates across the different states. We start considering the process of lumping two states of the model, A and B, into a single state X (Figure 3A). To preserve the out-going probability flux and have an identical equilibrium occupancy of the remaining states of the model, we impose that:

**Figure 3.**
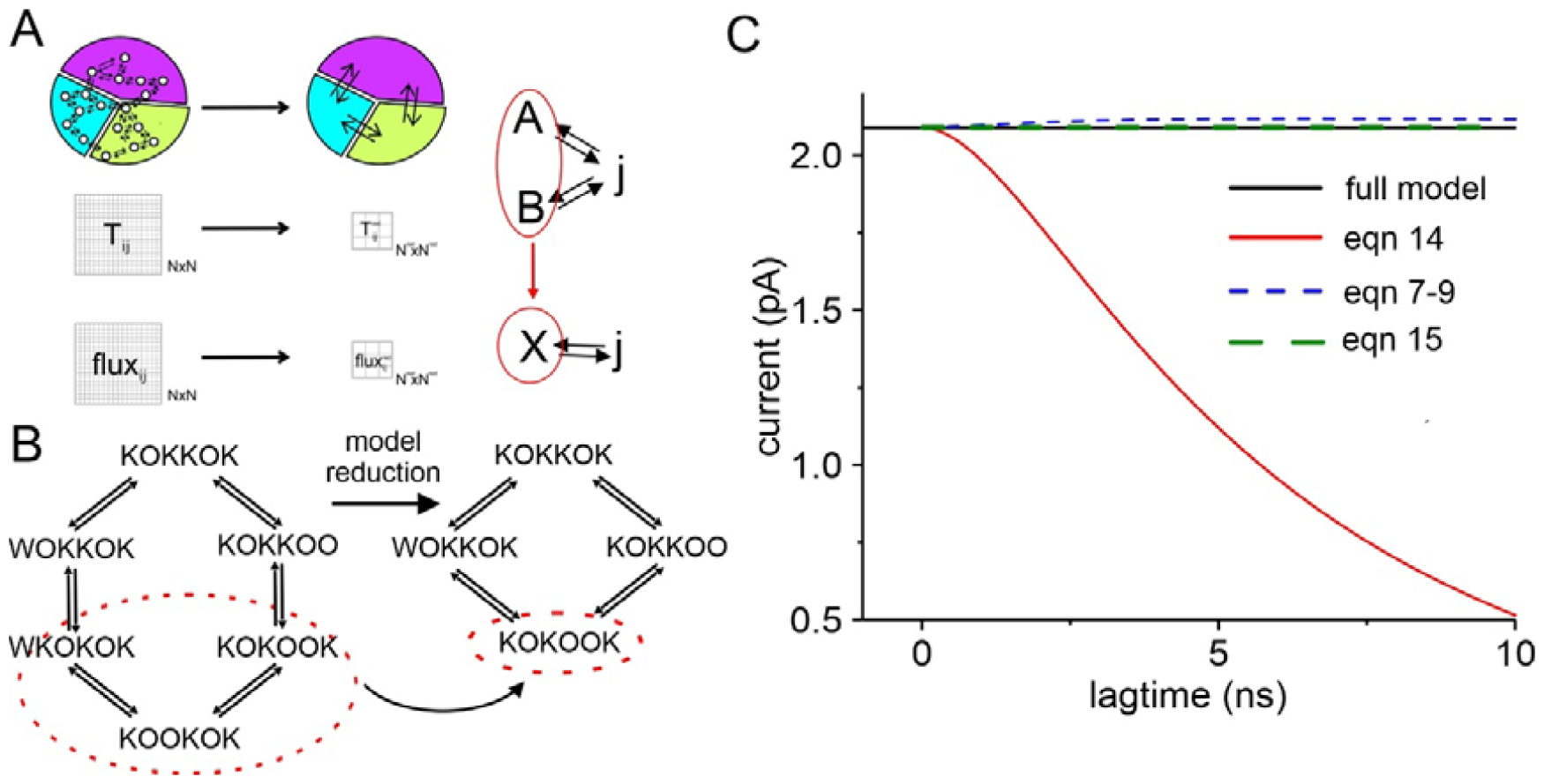
**A)** Drawing exemplifying the process of reduction of a MSM, considered for both the transition and the flux matrix, so to have a correct prediction of the ionic current. **B)** Scheme showing the reduction considered for the 6-state MSM taken as example. **C)** Ion current predicted for the reduced model using eqn 14 (red line), compared with the correct current of the full MSM (black line). The plot also shows the current recovered from transition probability matrices at varying lagtimes, using our proposed method of correction for large lagtimes, assuming (dashed blue) markovianity and assessing the transition matrices at the various lagtimes without assuming markovianity (green dashed line).

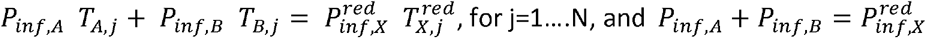

giving

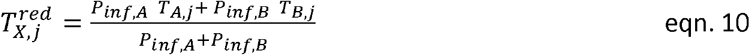

Similarly, to preserve the in-going probability flux we have:

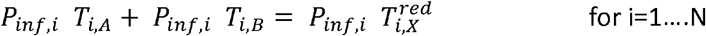

giving

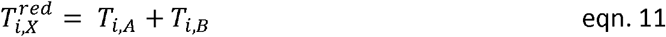

Eqns 10 and 11 can be applied multiple times to lump together as many states as desired. To assess the ion current using the reduced model, we also need to find a coherent reduced flux matrix. This can be done by imposing the in-going and out-going flux charge conservation in the following way:

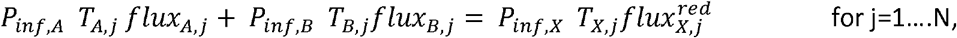

giving

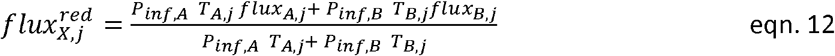

and

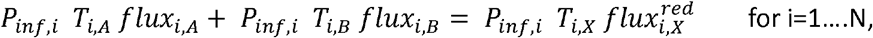

giving

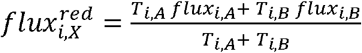

In the case of the term *flux*_*xx*_, we need to consider possible charge fluxes existing between the lumped states:

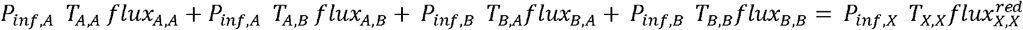

giving

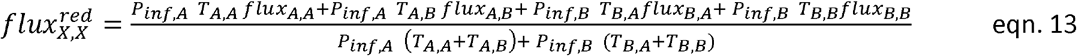

This reduction strategy is equivalent of starting from a count matrix containing all the transition events seen for the original N states and summing up all the events starting from or ending to the lumped states into a single value, as well as summing all the transitions in and out of the lumped states. The same result would be obtained starting from MD data by redefining the macrostate including all the state spaces of the two lumped states. Once the reduced transition probability matrix, T^red^ and the reduced flux matrix F^red^ are obtained, using eqns 10-13, the ion current of the reduced model can be predicted using eqn. 9:

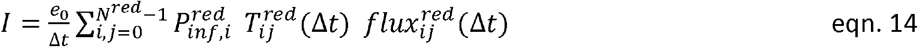

We will illustrate this reduction method using again the MSM considered in Figure 1B, in which states 3 (KOKOOK), 4 (KOOKOK), and 5 (WKOKOK) are lumped together (Figure 3B). Figure 3C plots the current predicted for the full MSM (eqn. 5, black line) and for the reduced, 4-state model (eqn. 14, red line). It is evident that the correct current is recovered, once again, provided that a sufficiently small lagtime is used for the determination of the transition probability matrix.

## The problem of non-markovianity

We showed that the ion current is correctly predicted only when very short lagtimes are used for constructing the transition probability matrices. Unfortunately, in the case of MD data, when lagtimes are that short, the transition probability matrices obtained are no longer markovian, in the sense that they cease to respect the Chapman-Kolmorokov equation (4). Thus, models constructed from MD data are actually quasi-MSM (qMSM), where markovianity is respected only for relatively large lagtimes. This occurrence obviously precludes the possibility of exploiting the very useful Markov properties on a very fast time scale, and this represents a significant limitation in the study of ion channel permeation, as shown later.

The non-markovianity in the low lagtime regime may be explained by the presence of multiple energy minima (microstates) within the states chosen to build the MSM. If the transitions between the microstates are not sufficiently fast compared to the chosen lagtime, the system can preserve memory of its previous (older than one lagtime) history, which makes it not-markovian. Also, the reduction procedure described above (Figure 3) to decrease the number of states of an MSM encounters the same problem. In fact, also in this case the new reduced model contains macrostates derived from the lumping of more energetically stable states, and this limits the markovianity at lagtimes sufficiently larger than the lifetime of the single states lumped together. In Figure 4, the black continuous lines represent the transition matrix elements (the element T_ij_ is represented by the plot in row *i* and column *j*) obtained for the reduced model, using eqn 10-11 on full model transition matrices taken at various lagtimes. Red and blue symbols in the plots represent instead the prediction of the Chapman-Kolmorogov equation (eqn.4) performed using transition matrices of the reduced model obtained for lagtimes of 1 and 20 ns, respectively. It is evident that significant deviations from the markovian prediction are present when the lagtime considered is 1 ns (arrows indicate the major discrepancies), while the predictions for a lagtime of 20 ns are essentially superimposed to the behavior of the reduced model. These results are in line with the previously exposed notion that the lumping of different microstates together generates a non-markovian behavior of the resulting reduced model, especially evident at short lagtimes.

**Figure 4.**
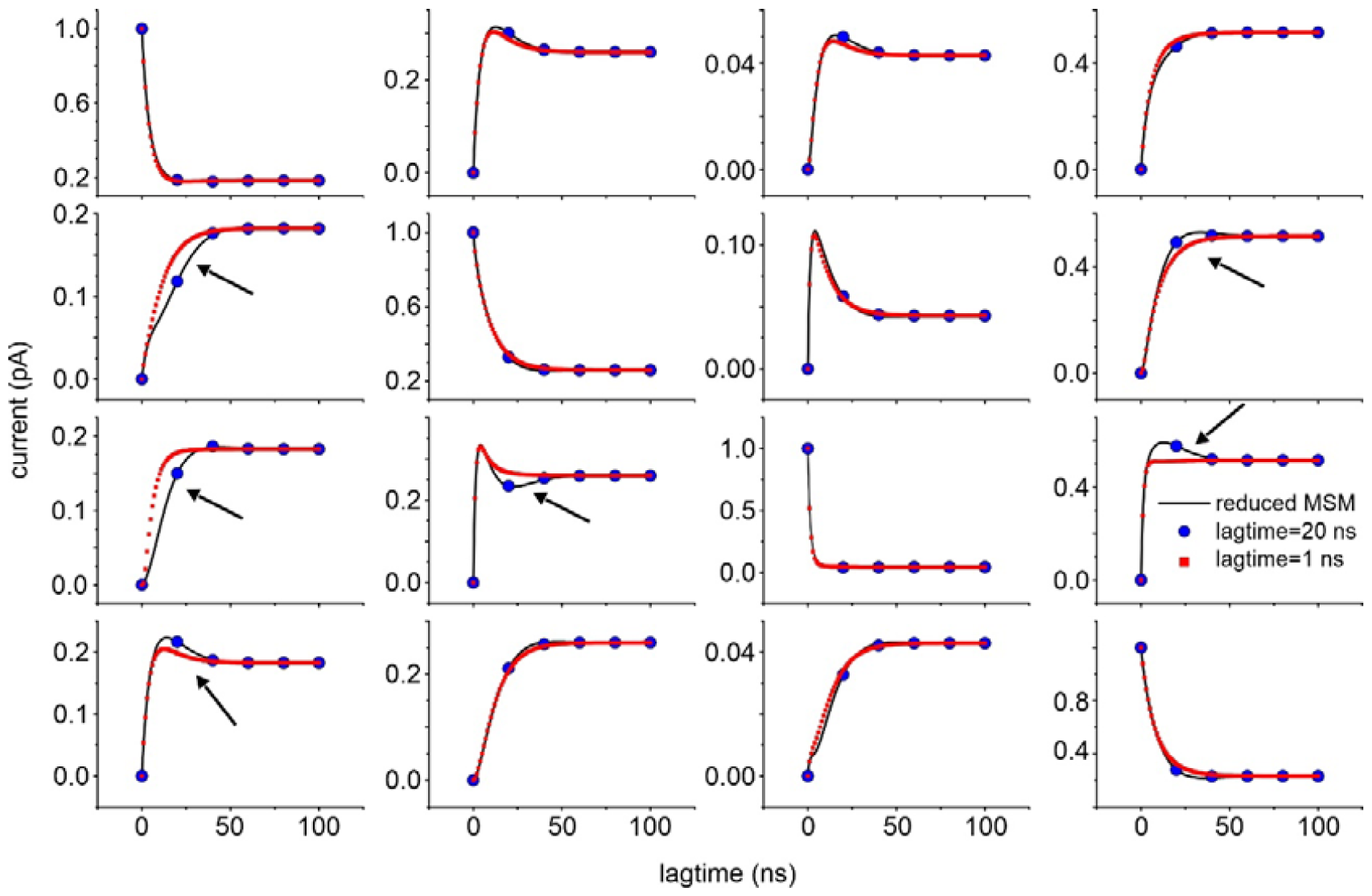
Transition matrix elements obtained for the reduced MSM (the plot in row i and column j represents the T_ij_ element of the matrix) vs lagtime (black lines) are compared with the Chapman-Kolmogorov prediction using a lagtime of either 1 (red symbols) or 20 ns (blue symbols).

The non-markovianity of a qMSM at short lagtimes represents a major limitation when trying to predict ion currents correctly. As we have seen in the previous paragraph, the prediction of an ion current can be done by first constructing a flux matrix valid only at very short lagtimes, and then progressively/recursively assessing composite flux matrices valid for larger lagtime transition matrices, by performing a weighted mean of fluxes resulting from all the possible pathways. However, it is important to note that, when using eqns 7 and 8 to recursively assess the composite flux matrix, we are assuming a markovian behavior of the system (eqn 7 is a for of the Chapman-Golmogorov equation), and thus this procedure will not produce a correct result in the case of a reduced (quasi-Markov state) model. This is demonstrated in Figure 3C, where the blue dashed line, obtained using eqns 7-9 that assumes a markovian behavior, slightly but significantly deviates from the correct value of the current. A possible solution to this problem is to consider the transition probability matrices directly obtained at various lagtimes, instead of predicting them by assuming a markovian behavior and using eqn. 7. More specifically, we propose the use of the following equation to recursively assess the composite flux matrix:

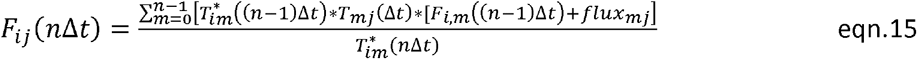

Where T^*^ represents transition probability matrices assessed directly from MD data at varying lagtimes, instead of using eqn. 7. The green dashed line in Figure 3C, representing an ion current prediction performed using transition probability matrices directly obtained for the reduced model and performing the recursion with equation 15, demonstrates that this procedure recovers the correct current at all lagtimes.

### Assessing the rate constants of the MSM of ion permeation

The methods described above allow, starting from MD data, to obtain an MSM for ion permeation containing relatively few states, with a transition probability matrix and associated flux matrix at lagtimes at which the model is fully markovian. The next step is to estimate the Q matrix containing the rate constants associated to each possible transition present in the model. This can be done using a fitting procedure that attempts to find the Q matrix that minimizes the following function:

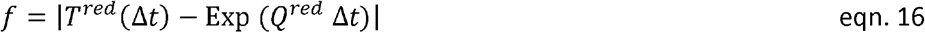

Where *T*_*red*_ (Δ *t*) is the transition matrix of the reduced model, obtained as described at the lagtime Δ *t*, and *Q*_*red*_ contains the rate constants to be found. In our reduced model, with a lagtime of 20 ns and using the minimization strategy described by eqn 16 and eqn 15, we obtained the following Q and F matrices:

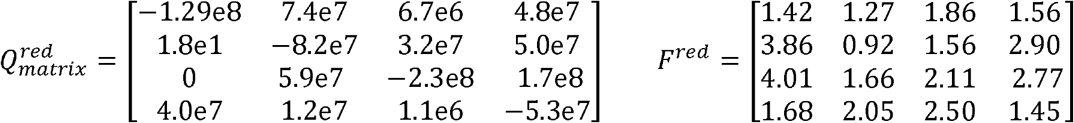

from which a predicted current of 1.98 pA is obtained, using eqn. 14, which is quite close to the real value of 2.089 pA. Notice that the minimization of eqn 16, based on the definition of the generator matrix given in eqn 2, recovers the correct Q matrix only for a fully markovian MSM. This implies that in presence of a quasi-MSM we need to perform the minimization using transition probability matrices obtained at lagtimes sufficiently large to guarantee a markovian behavior. This is shown in Figure 5 that reports the current predicted using a Q matrix obtained from transition probability matrices at varying lagtimes. It is evident that a correct prediction requires lagtimes close to 20 ns, that guarantee a markovian behavior (cf Figure 4C).

**Figure 5.**
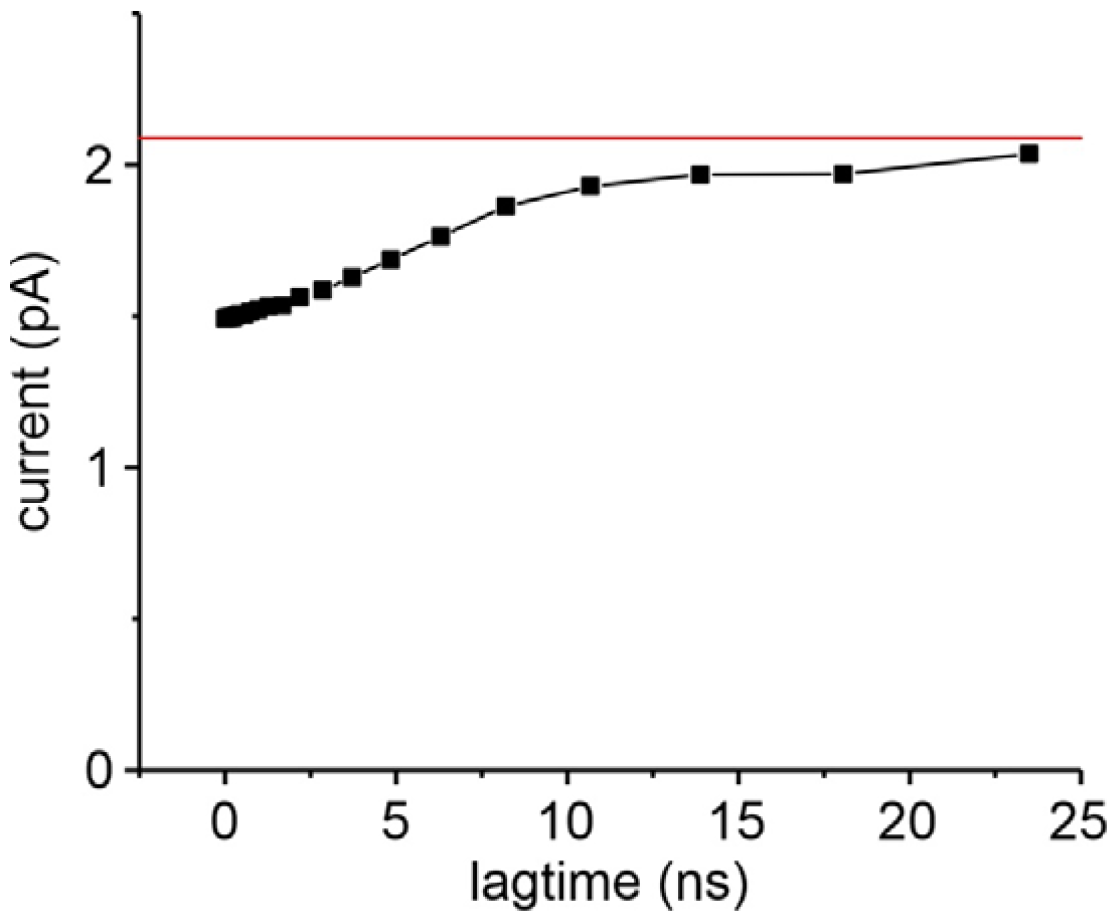
Currents estimated using eqn 5, with a Q matrix recovered from fits (eqn. 16) where reduced transition probability matrices at varying lagtimes were used. It is evident that a correct current is predicted only when transition probability matrix obtained at large lagtimes are used in the minimization procedure. The red line represents the correct value of the current for the considered model.

### Overall suggested algorithm

To summarize, we propose the following procedure to extract a reduced MSM from MD data:

1) Obtain transition matrices for ion permeation at varying lagtimes from MD data.

2) Assess the full model flux matrix associated to the shortest lagtime transition matrix available, using the algorithm in appendix B. The use of a sufficiently short lagtime can be verified by comparing the current predicted by the transition and flux matrices (cf eqn 6) with the ion current directly assessed by MD simulations. A predicted current significantly lower than the one assessed from MD is indicative of a too large lagtime used.

3) Reduce the model at short lagtimes by lumping together states and finding reduced transition probability and flux matrices through eqns 10-13. The correctness of the reduction procedure can be verified by monitoring the current predicted by the reduced model at the short lagtime using eqn 14. Different strategies of model reduction can be considered, based on the selection of the most physically significant states, on the relevant state occupancy, and other parameters.

4) Test for markovianity of the transition matrix. If it fails, increase the lagtime until finding a nearly markovian behavior. Assess flux matrices at large lagtimes, at which the model is fully markovian, using eqn 15. This will allow using the model for predicting the ion current from the transition probability matrix obtained at large lagtimes.

5) Assess the rate constants associated to each transition of the reduced model using a fitting procedure that minimizes eqn 16, and using a fully markovian transition probability matrix, obtained at large lagtimes.

The correctness of the procedure can be monitored by assessing at the various stages the predicted ionic current.

## Results

We used the above algorithm to find a reduced model of K permeation through the KcsA channel from transition matrices built at various lagtimes using MD data at +200 mV of applied potential. We performed a total of 29.69 µs of simulation time in 8 replicas, obtaining 334 permeation events, resulting in a current of 1.79 pA. As reported in Domene et al., 2021 (7), the transition matrix was determined by looking at the presence of K and/or water within the six different sites S0 to S5 (going from the extracellular side to the intracellular cavity). Of the whole set of 4096 possible configurations, only 122 were actually present in the original transition matrix, i.e., had been effectively found in the MD simulation. Figure 6A shows the occupancy of the various filter configurations at steady state (only the configurations with an equilibrium occupancy higher than 0.1% are shown). The result is that the most stable configurations are those having a K ion at sites 0, 2, 3, and 5, with sites 2 and 3 always occupied, regardless of the presence of water in the various sites.

**Figure 6.**
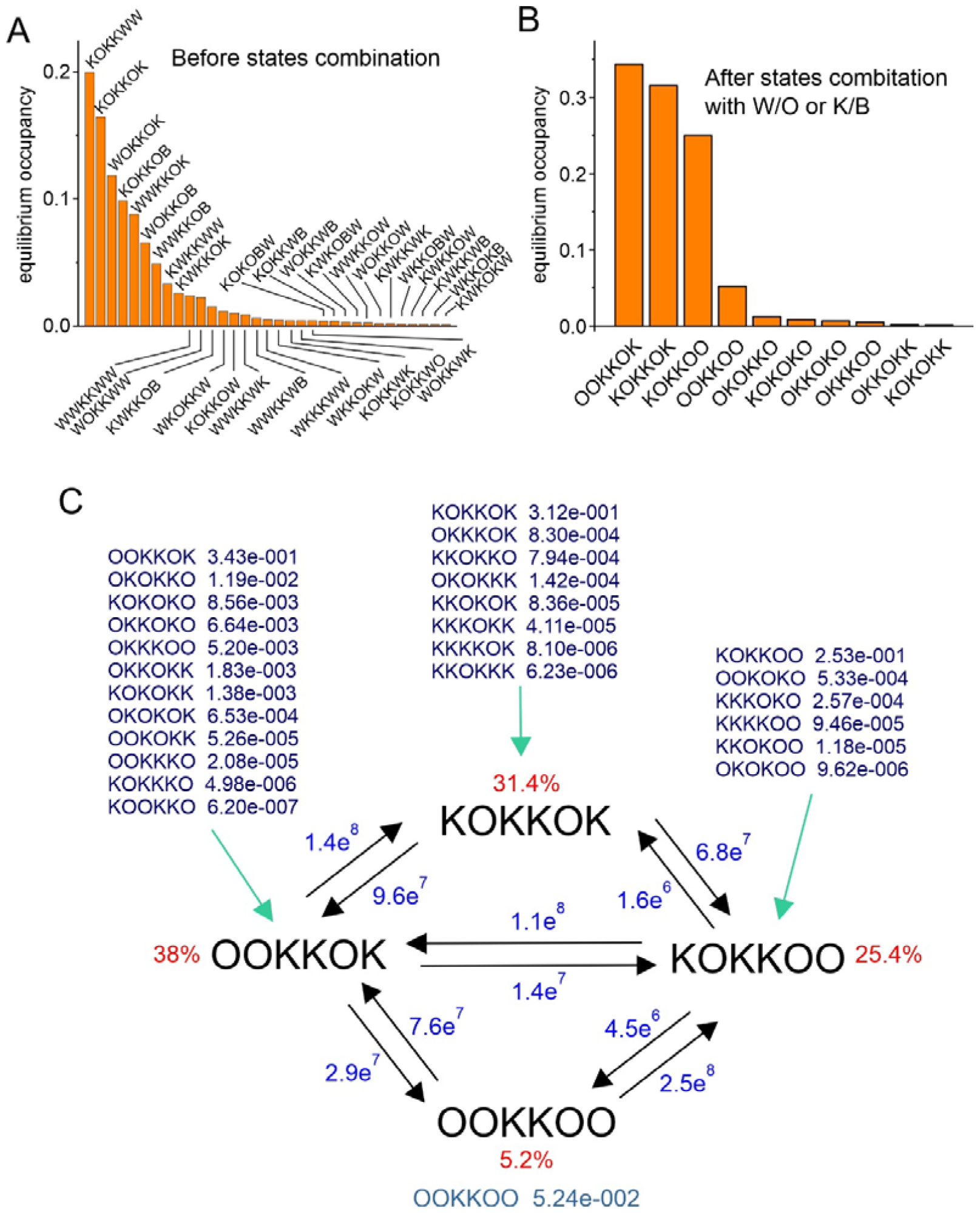
**A and B)** Plots of the equilibrium occupancy of the various state configurations found in the MD data, before (A) and after (B) lumping together states having the same K ions asset regardless of the presence of water in the binding sites. In both cases only states with an occupancy higher than 0.1% are shown. **C)** Reduced kinetic scheme obtained for the MD data at 200 mV of applied potential. In black are shown the states and relative occupancy contributing to each reduced state, in blue the rate constants obtained by the minimization procedure, and in red the relative steady state occupancy.

Since we are mainly interested in the permeation of K, we performed a combination of states that had the same K ion configuration. As an example, states K0KK0K, K0KK0B, K0KKWK, K0KKWB, KOKBOK, KOKBOB, etc, were all combined in one configuration indicated as K0KK0K, where this time K indicates the presence and 0 indicates the absence of a K ion in that site, regardless of the presence of water. As we have shown above, state combination was performed using a method that preserves the steady-state occupancy of the remaining states, as well as ingoing and outgoing ion fluxes. In addition, it has been clearly shown that the permeation process does not involve water transport, since water molecules are totally excluded from the internal S2 and S3 sites (12). The described combination allowed to further reduce the number of states from 122 to 27, as shown in decreasing order of steady-state occupancy in Figure 6B. Once again, it is possible to notice that the most represented states are those having sites S0, S2, S3, and S5 occupied by K, while only rarely sites S1 and S4 bind the permeating ion.

Finally, we performed a further combination of ion configurations, using the following criterion: if a configuration *i* was connected to a configuration *j* with a P*ij*/P_*ji*_ ratio relatively large, meaning that the *i* to *j* transition could be considered almost irreversible, then the *i* and *j* configurations were combined using the same method described above, and the resulting combined configuration was named as the configuration of the absorbing state *j*. Using a P*ij*/P_*ji*_ minimum ratio of 13 (a value that guarantee that states with an occupancy over 1% remained separated) as combination criterion, we reached a reduced model of permeation including only 4 states, namely OOKKOO, OOKKOK, KOKKOO, and KOKKOK. The transition matrix and corresponding flux matrix of the reduced model, both assessed for a lagtime of 0.01 ns, that corresponded to the frequency of stored trajectories of the MD simulation, are:

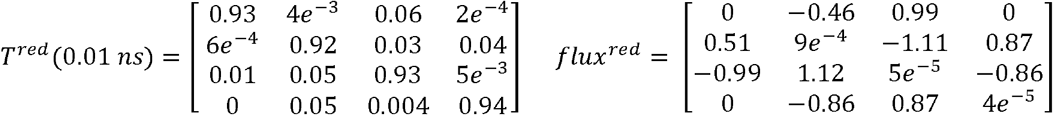

Figure 6C shows the kinetic scheme of the reduced model obtained, together with the states contributing to each of the macrostates of the model (with the relative occupancy). Notice that this reduced model is not perfectly markovian because of the very short lagtime (0.01 ns) used to define transition and flux matrices.

Once obtained a reduced qMSM model of K permeation through the KcsA channel, we estimated the transition probability matrix and corresponding flux matrix at higher lagtimes, using the method described in the previous section, to have a fully-markovian model of K permeation. Figure 7 illustrates transition matrix derived from MD data at 200 mV and the prediction of the Chapman-Kolmogorov equation at different lagtimes. The black continuous lines represent the transition matrix elements (the element T_ij_ is represented by the plot in row *i* and column *j*) obtained for the reduced model as a function of the lagtime. Red and blue symbols in the plots represent instead the prediction of the Chapman-Kolmogorov equation performed using transition matrices of the reduced model obtained for lagtimes of 0.2 and 4 ns, respectively. It is evident that significant deviations of the markovian predictions from the true transition matrices are present at 0.2 ns lagtime, while the predictions for a lagtime of 4 ns are closer to the real behavior, although differences can still be appreciated. We thus decided to assess the reduced transition probability matrix and flux matrix for a lagtime of 10 ns, at which we expected an essentially markovian behavior. The transition matrix and flux matrix estimated at 10 ns lagtime were:

**Figure 7.**
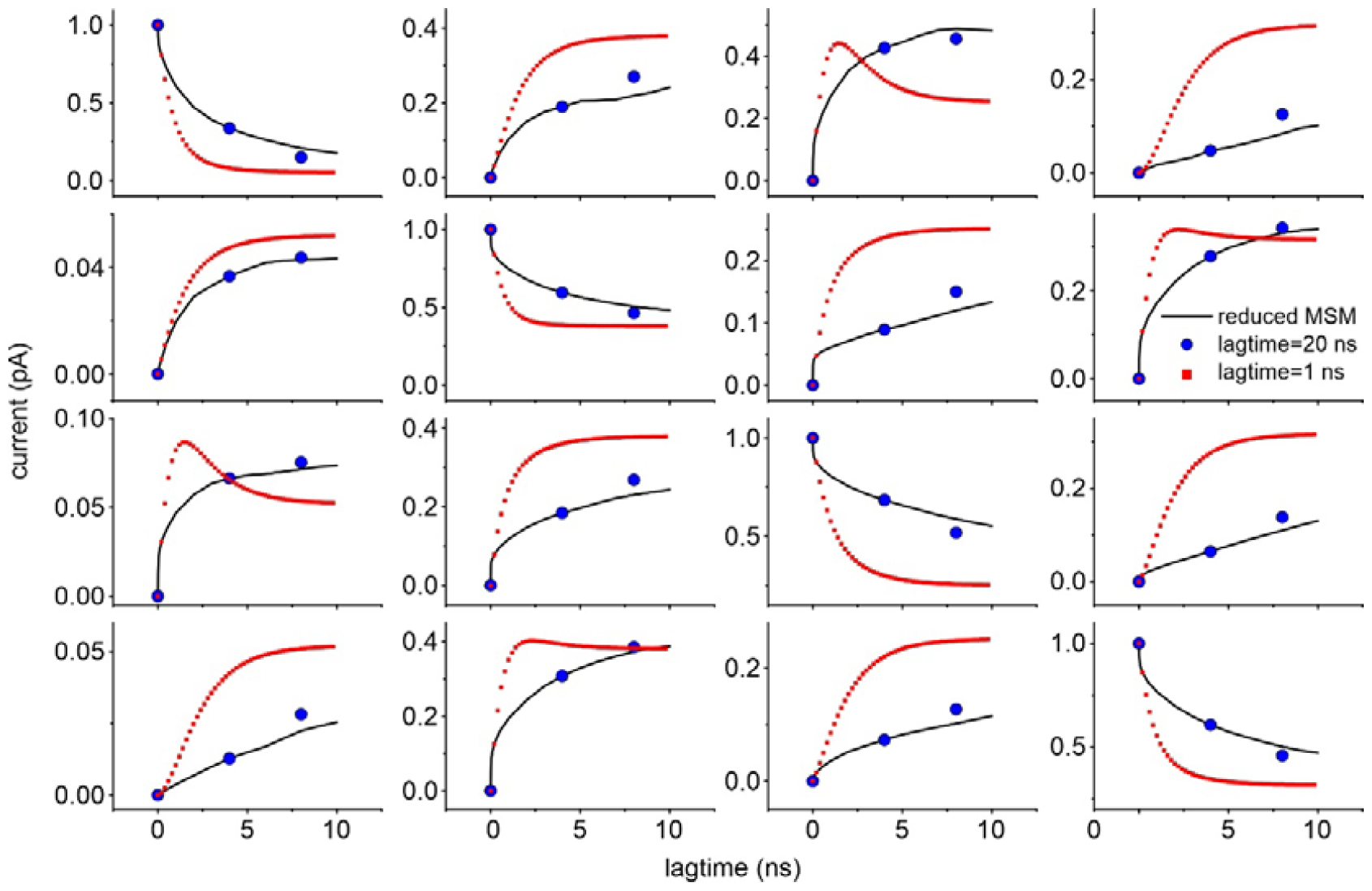
Transition matrix obtained for the MD data at 200 mV (the plot in row i and column j represent the T_ij_ element of the matrix) vs the lagtime (black lines). Data are compared with the Chapman-Kolmogorov prediction using a lagtime of either 0.2 (red symbols) or 4 ns (blue symbols).

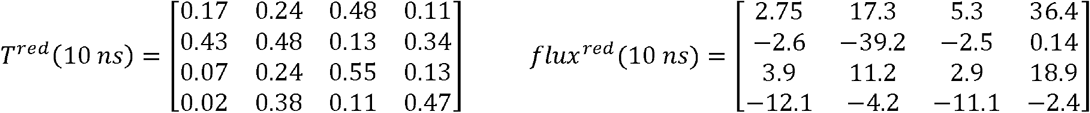

As shown in Figure 8, our method of assessment of the composite flux matrix at higher lagtimes predicts a current similar to that directly obtained from MD at all lagtimes, suggesting that the method used is robust.

**Figure 8.**
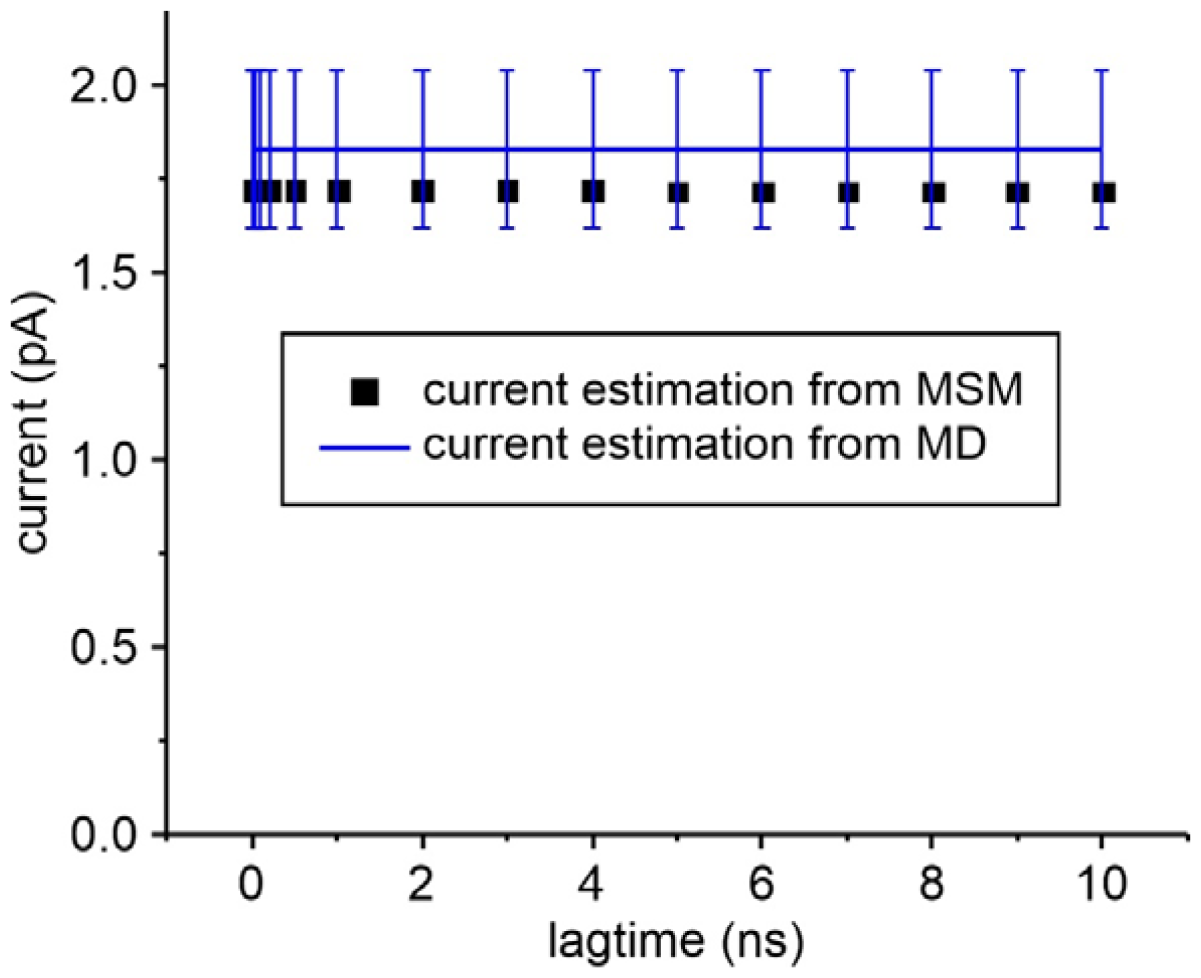
Predicted current using transition probability matrices and composite flux matrices at varying lagtimes. The blue line and errors represent the mean and standard error of the ionic current assessed in the 8 MD replicas.

Finally, we performed a fit of the transition matrix obtained for a lagtime of 10 ns to obtain the transition rate constants to be associated to the MSM of K permeation. The blue numbers in the scheme of Figure 6C report the best fit parameters obtained. Based on their values, the transitions OOKKOO->KOKKOK and KOKKOK->OOKKOO are essentially absent (their values were below 0.1 s^-1^, and thus the corresponding arrows are not shown in the model), as expected from the fact that they involve the movement of two K ions simultaneously. These results allow us to propose a model identical to the association-dissociation model previously proposed (13, 15). In this model K permeation can be viewed as a K ion binding to either one of the two external sites, S5 or S0, and another K ion being released (unbinding) from the opposite site (upper part of the scheme). In fact, the two events may also occur in the reverse order, with first the unbinding of a K ion and then the binding of another K ion at the opposite site (lower part of the scheme). For the system to give a continuous K flux we find that the two 3-K ion configurations, KOKKOO and OOKKOK, can interconvert one into another through a single file movement.

## Conclusions

The method reported in this paper combines MD simulations and MSM with the aim of allowing advancements in the field of ion channel permeation. More specifically, using the method we have described, it is possible to build a reduced MSM of ion channel permeation characterized by a relatively low number of states, and able to quantitatively predict the correct ionic current. The model can be used mainly for two distinct purposes, to compare MD results to experiments and to understand the mechanism of ion channel permeation in terms of the most important ionic movements inside the ion channel pore.

We applied the method to MD simulation data obtained at 200 mV, in presence of 200 mM K+, for one of the most studied K channels, the KcsA, and found not only that it is able to reproduce the amplitude of currents, but to suggest a very specific K permeation mechanism. Following the state reduction procedure described, we obtained an MSM with only 4 main states, and transition and flux matrices capable to predict essentially the same ionic current directly measured in the MD simulation, by counting the number of K ions passing through the pore. The results of the analysis also suggest that ion permeation in KcsA channels follows an association-dissociation scheme, meaning that the binding/unbinding of a K ion at one entrance of the channel promotes the unbinding/binding of a K ion at the other channel entrance. The two processes, combined with the ability of K ions inside the selectivity filter to move in single file, generate a cyclic process promoting the continuous passage of permeant ions from one side of the membrane to the other.

As we have seen, a major advantage of the proposed method is to link the atomic structure of the ion channel pore to the main experimental data in ion channel permeation, i.e., the single channel current. Although the currently available resources are sufficient to estimate ion currents using MD simulation, the high computational cost limits the range of analyses and imposes the use of extreme boundary conditions in terms of membrane potentials and ion concentrations. These limitations impact most strongly when one wants to study the functional properties of an ion channel for which the shape of the current-voltage relationship is much more informative than a current estimate at a single membrane potential. In this case, it would be necessary to simulate several membrane potentials, including some close to equilibrium conditions. Unfortunately, the computational cost of MD simulations prevents such investigations today, so we suggest to get around this problem by constructing MSMs of ion permeation to predict I-V relationships under conditions and voltages not directly tested with MD, thereby optimizing computational resources in ion permeation. More specifically, by applying our method to a limited number of MD simulations at high membrane potentials, it might be possible to determine the voltage dependence of the various kinetic rate constants of the reduced MSM. This would make it possible to predict the ion current even at more physiological voltages not directly tested in MD and the entire I-V relationship for the single-channel current (Furini et al., Ms in preparation).

The application of our proposed method would in addition greatly help understand the main determinants of the permeation mechanisms in a variety of ion channels. Currently, MD provides such an overwhelming amount of data that makes it extremely difficult to identify the main determinants of the permeation process. The approach of analyzing MD data and obtaining MSMs of ion channel permeation, and then repeatedly applying the state reduction technique proposed in this paper, would allow the identification of a few relevant metastable states, and a few transitions that primarily define the permeation process. Most importantly, during the second half of the last century many relatively simple kinetic schemes, which are essentially reduced Markov models, were proposed to explain the permeation and selectivity of many different ion channels (16–19). These simple mechanisms have been extensively tested for their ability to accurately reproduce many experimental results. The approach proposed in this paper could be used, once the atomic structures of those channels have been determined, to understand in terms of single aminoacidic residues and atomic structure, the significance of the states and transitions present in each of these kinetic models. Above all, the permeation process is reduced to few states/transitions and becomes more easily comprehensible.

## Acknowledgments

This work has been supported by the project “Kinetic models of ion channels: from atomic structures to membrane currents” funded by MUR PRIN 2022 (grant 20223XZ5ER, CUP J53D23006940006). SF acknowledges CINECA for awarding access to computational resources through the ISCRA initiative (grant number HP10BJPCFW).

## Appendix

### Appendix A Relationship between transition probability matrix and Q matrix for a Continuous time Markov process

A stochastic process Z(t), defined over a discrete state space S of cardinality N is a Continuous time Markov process (CTMP) if for any sequence of times (t_0_<t_1_…….<t_m-1_<t_m_),

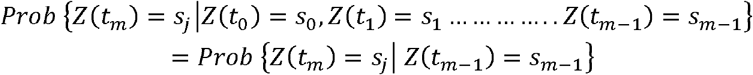

Let’s call *Prob* {*Z* (*t*_*m*_) = *s*_*j*_ ∣ *Z*(*t*_*m*−1_} = *T*_*ij*_ (*t*_*m*−1_, *t*_*m*_), elements of the matrix of conditional probabilities T(t_m-1_,t_m_), and *Prob* {*Z*(*t*) = *s*_*j*_} = *p*_*i*_ (*t*) row vector of occupancies p(t).

The Chapman-Kolmorogov (C-K) equations implies that:

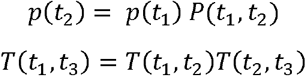

A CTMP is time homogeneous if the conditional probability T(t_1_,t_3_) depends only on the difference t_3_-t_1_ T(t_1_,t_3_)= T(t_1_+ t_2_,t_3_+ t_2_)=T(t_3_-t_1_). In this case the C-K equation becomes

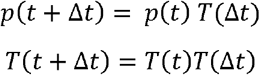

Let’s define the Q matrix with elements 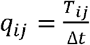 for small Δ*t*, and *q*_*ii*_ = − Σ*j q*_*ij*_

If follows that *T*_*ij*_ (Δ*t*) = *q*_*ij*_ Δ*t* and *T*_*ii*_ (Δ*t*) = 1 + *q*_*ii*_ Δ*t*

The homogeneous C-K equation then becomes:

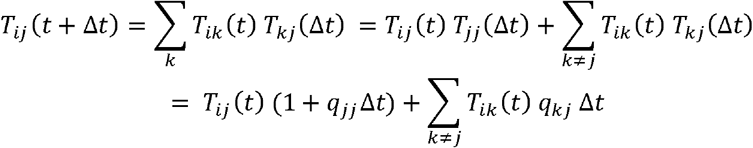

It follows

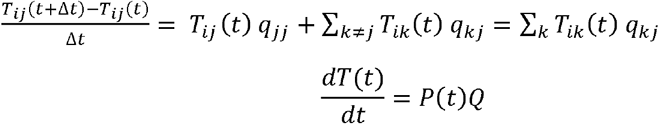

The above equation has the formal solution

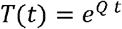

relating the transition probability matrix to the Q matrix

### Appendix B Flux matrix assessment

1. Starting from an initial configuration i other configurations are produced by allowing the entrance of water or K ions to the external sites S5 and S0, or moving water and K ions from a binding site to the adjacent one inside the SF to the left or to the right.
2. For each new configuration obtained, an associated flux is assessed by adding +1/7 for a K movement towards the external side, and -1/7 for each K movement towards the internal side. For example, a flux of 1/7 is assigned to the transition KOKKOO → KOKOKO, since the configuration KOKOKO is obtained by moving one step to the right the K ion in the S2 site.
3. Point 2 is repeated until the configuration j is reached, and the i → j flux is assessed by summing all the fluxes associated to the intermediate transitions while going from i to j. For example, a flux of +2/7 is associated to the transition KOKKOO → KOKOOK if this transition is obtained by the two consecutive transitions KOKKOO → KOKOKO and KOKOKO → KOKOOK both having associated a flux of +1/7.

## Notes

### Competing Interest Statement

The authors have declared no competing interest.

